# Hunger improves reinforcement-driven but not planned action

**DOI:** 10.1101/2021.03.24.436435

**Authors:** Maaike M.H. van Swieten, Rafal Bogacz, Sanjay G. Manohar

## Abstract

Human decisions can be reflexive or planned, being governed respectively by model-free and model-based learning systems. These two systems might differ in their responsiveness to our needs. Hunger drives us to specifically seek food rewards, but here we ask whether it might have more general effects on these two decision systems. On one hand, the model-based system is often considered flexible and context-sensitive, and might therefore be modulated by metabolic needs. On the other hand, the model-free system’s primitive reinforcement mechanisms may have closer ties to biological drives. Here, we tested participants on a well-established two-stage sequential decision-making task that dissociates the contribution of model-based and model-free control. Hunger enhanced overall performance by increasing model-free control, without affecting model-based control. These results demonstrate a generalised effect of hunger on decision-making that enhances reliance on primitive reinforcement learning, which in some situations translates into adaptive benefits.

**Significance statement:** The prevalence of obesity and eating disorder is steadily increasing. To counteract problems related to eating, people need to make rational decisions. However, appetite may switch us to a different decision mode, making it harder to achieve long-term goals. Here we show that planned and reinforcement-driven actions are differentially sensitive to hunger. Hunger specifically affected reinforcement-driven actions, and did not affect the planning of actions. Our data shows that people behave differently when they are hungry. We also provide a computational model of how the behavioural changes might arise.

## Introduction

Hunger is an adaptive motivational state that drives us to eat, restoring homeostatic balance (Saper et al., 2002). However, maladaptive behaviour in the context of hunger generates many clinical problems. One in 7 adults suffer from obesity, one of the most serious health issues in the developed world, and 10% suffer from a range of eating disorders. One critical contributing factor may be the changes in decision-making produced by hunger.

One well-studied effect of hunger is to amplify the value of reward, both specifically for food (Epstein et al., 2003; Malik et al., 2008), and even for generic reward signals (Aitken et al., 2016; Cone et al., 2016; Haase et al., 2009; Papageorgiou et al., 2016; Siep et al., 2009) (Briers et al., 2006; Simon et al., 2017). This modulation of value may feeds into primitive reinforcement systems (Daw and O’Doherty in Glimcher and Fehr (2013)), strengthening simple action learning. This kind of value-based action reinforcement is inflexible, and is more strongly engaged in people with eating disorders (Voon et al., 2015). However, to allow for flexible, context-sensitive action selection, these primitive systems must be guided by a planning system, which computes *how* to get to a reward using a causal model of the world, rather than simply engaging actions that previously led to reward. The planning system is sensitive to metabolic needs insofar as they determine our goals (Aw et al., 2009; Dickinson and Balleine, 1994; Dickinson, 1985; Friedel et al., 2014), but it is not known whether needs can make us generally better or worse at goal-directed planning.

In this study, we asked whether hunger promotes primitive decisions based on reinforcement, or planned actions. Simple reinforcement is model-free, with memory only about how rewarding actions were in particular states. In contrast, the model-based planning system, uses knowledge about how actions change the state of the world. Would both of these systems be sensitive to biological needs?

On one hand, we might expect the model-free system to be modulated by hunger, because it may rely more on primitive neural systems. Circulating hormones that signal metabolic need target subcortical brain areas that regulate feeding (Elmquist et al., 1998; Zigman et al., 2006). In particular, leptin inhibits and ghrelin activates dopaminergic neurons in the ventral tegmental area, and could therefore modulate learning and decision-making via the mesolimbic pathway (Abizaid et al., 2006; Figlewicz et al., 2007; Hommel et al., 2006). This automatic mechanism could presumably be advantageous, since hunger may trigger reward-driven behaviour enabling organisms to adapt to the variability in environmental abundance (de Ridder et al., 2014; Levy et al., 2013; Shabat-Simon et al., 2018; Symmonds et al., 2010).

On the other hand, we might expect a higher-level, model-based system to be modulated by hunger. It is by definition more flexible, being sensitive to state and context. Higher-level control may be decreased by hunger. For example, hunger promotes impulsivity, preventing us from reaching long-term goals (Bartholdy et al., 2016; Kirk and Logue, 1997; Skrynka and Vincent, 2019). Cognitive control is regulated by homeostatic hormones (Higgs et al., 2017), and is impaired in obesity (Hassenstab et al., 2012) but enhanced in anorexia nervosa (Compan et al., 2015).

In the present study, we employed a two-stage decision-making task (Daw et al., 2011) to examine if hunger modulated the model-free or model-based decision system. In this task, people must choose between two rockets, each going predominantly to one of the second-stage planets (Fig 1A). Once on a planet, they choose between two aliens that probabilistically deliver a reward. If a reward is obtained, it pays off to return to that planet on the next trial by selecting the same first-stage rocket again. Crucially, the rockets occasionally go to the other planet (a “rare transition”). If a reward is earned in this situation, it pays off to choose the *other* rocket next time, because that rocket is more likely to bring you back to the same second-stage planet. The simple, “model-free” approach is to repeat the rocket choice whenever it was rewarded. A more sophisticated, “model-based” strategy is to consider that rockets do not always go to the same planet and plan ahead to reach the most rewarding planet.

**Figure 1.**
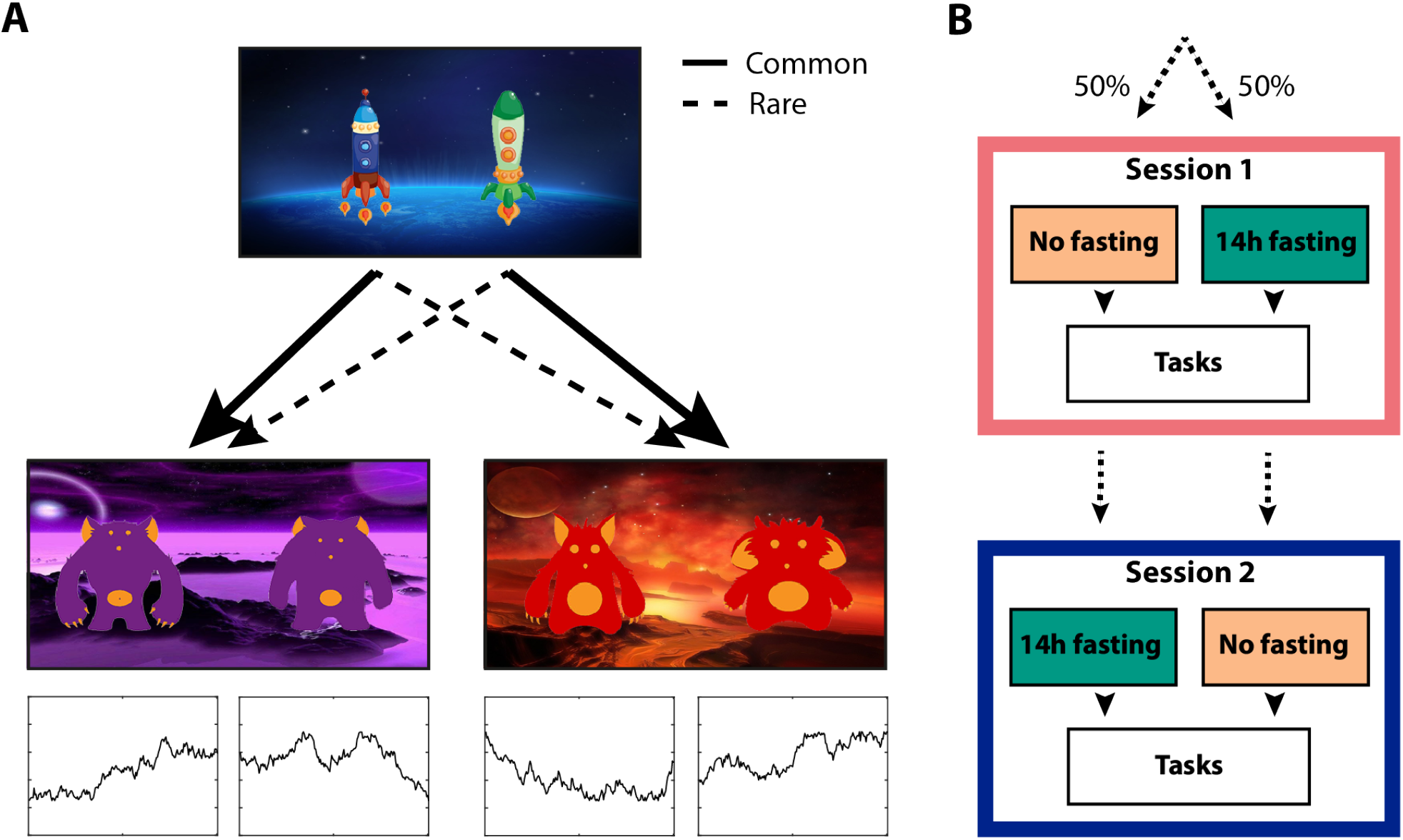
Study design. **A)** Schematic of the task design developed by (Daw et al., 2011). Each first-stage choice rocket flew predominantly to one of the second-stage planets (common transition: 70% of the trials) and sometimes to the other second-stage planet (rare transition: 30% of the trials). Each second-stage alien probabilistically led to a reward. The reward probabilities for each second-stage alien fluctuated across trials between 25% and 75% according to a Gaussian random walk. Task instructions and images were obtained from (Kool et al., 2016). **B)** Participants were tested in a within-subjects counterbalanced, randomised crossover design. Participants were tested on two separate days approximately 1 week apart. One session took place after 14 hours fasting, the other session after consuming a full meal.

We showed that increased metabolic need enhanced reflexive model-free control of behaviour, without affecting deliberate model-based control. Although food deprivation has been shown to increase impulsivity, this study provides evidence that relying on a reflexive model-free system does not make hungry people maladaptive, when the deliberate model-based system remains unaffected.

## Methods

### Participants

Thirty-two healthy volunteers (females: 20, mean age: 25.6 ± 6.5) were recruited for this study. All participants were healthy, had no history of psychiatric diagnoses, neurological or metabolic illnesses, and had not used recreational drugs in the past 3 months. All participants had a normal weight (Body Mass Index: 22.9 ± 3.2 kg/m^2^), regular eating patterns and no history of eating disorders. Each participant gave written informed consent and the study was conducted in accordance with the guidelines of the University of Oxford ethics committee.

### Manipulation of metabolic state

Participants were tested in a within-subjects counterbalanced, randomised crossover design for the effects of food deprivation on planning (Fig. 1B). Sessions were approximately 1 week apart (at least 4 days, but no more than 14 days). All sessions took place at the same time of day between 10 am and 1 pm, to minimise time-of-day effects. For one session, participants were asked to refrain from eating and drinking caloric drinks from 8 pm the night prior to testing. For the other session, participants were asked to eat normally the day before and consume a full breakfast within 1 hour of arriving at the lab for testing. We assessed the effect of food deprivation on self-reported feelings of hunger and mental state using a computerised Visual Analogue Scale (Bond and Lader, 1974; Flint et al., 2000).

Participants were asked to place a cursor on a 100 mm scale with positive or negative text ratings anchored at either end. This assessment provided a subjective measure of whether the manipulation worked. Participants also performed a risk-taking, attention and learning task not described in this paper.

### Paradigm

In the two-step task, participants make a series of choices between two stimuli, which lead probabilistically to one of two second-stage states (Fig. 1A). Each first-stage rocket leads more frequently (70%) to one of the second-stage states (a “common” transition), and in a minority of the choices (30%) to the other second-stage state (a “rare” transition). These second-stage states require a choice between two aliens that offer different probabilities of obtaining a monetary reward. To encourage learning, the reward probabilities for each second-stage alien fluctuated across trials between 25% and 75% according to a Gaussian random walk (Daw et al., 2011).

To solve this task, a model-free strategy would involve choosing the same rocket when it previously resulted in a reward. This would occur irrespective of whether the transition to the second-stage was common or rare (i.e. whether the planet was expected or unexpected), because the model-free system is insensitive to the structure of state transitions within the task. In contrast, a model-based strategy would necessitate *switching* to the other rocket after a reward, if the transition was a rare one (i.e. if the chosen rocket went to the unexpected planet). This is because participants can use their model of which rocket tends to lead to which planet, to maximise their chances of obtaining future rewards.

The task consisted of 201 trials, divided into three blocks of 67 trials separated by short breaks. If participants failed to enter a choice for a first or second-stage choice within 2 seconds, the trial was aborted and not included in further analyses. A new set of randomly drifting reward distributions was generated for each participant. Participants received 1 pence for every point they earned.

Before completing the full task, participants were extensively trained on different aspects of the task. Participants sampled 25 times from aliens with different reward probabilities. They were also told that each rocket preferentially goes to one of the planets, but they were not instructed on which rocket goes to what planet or the explicit transition probabilities (Kool et al., 2016). Finally, participants practised the full task for 25 trials. There was no response deadline for any of the sections of the training phase. Different rockets, planet colours and aliens were used for the training and experimental phase, and these were counterbalanced across conditions and participants.

### Stay-Switch Behaviour

The one-trial-back stay-switch analysis is the most widely used method for characterising behaviour on the two-step task. This method quantifies the tendency of a participant to repeat the choice made on the last trial or switch to the other choice, as a function of the outcome and transition on the previous trial. We considered four possible outcomes: Common-Rewarded (*CR*), Rare-Rewarded (*RR*), Common-Unrewarded (*CU*), and Rare-Unrewarded (*RU*). Model-based and model-free indices were computed from the stay probabilities following each outcome according to:

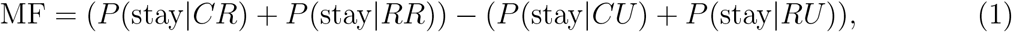

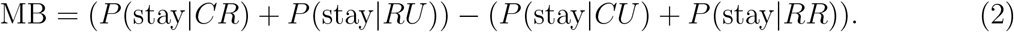

We also examined whether hunger modulated other measures of simple reinforcement learning. We found no effect of hunger on changes in model-free control for second-stage choices or for action-specific, stimulus-irrelevant choices at the first-stage (Fig. S1).

Statistical analyses were implemented in MATLAB and SPSS (IBM Corp. Released 2019. IBM SPSS statistics for Windows, Version 26.0. Armonk, NY: IBM Corp.).

### Computational modelling

We used a hybrid computational model that assumes that the model-based and model-free systems both contribute to choice behaviour (Daw et al., 2011). Model-based and model-free algorithms learn the value of the stimuli that appear in the task in three different pairs. There is one first-stage pair (*s*1 ∈ {1, 2}) and two second-stage pairs consisting of two stimuli each (*s*2 ∈ {3, 4, 5, 6}). The indices *s*1 and *s*2 refer to stage 1 and stage 2, respectively. The index *t* refers to the trial number.

At the first-stage, model-free ‘cached’ values were updated using a temporal difference algorithm. This algorithm learns to maximise the total outcome by strengthening or weakening associations between the first-stage state and the first-stage actions, depending on whether a reward followed that action or not:

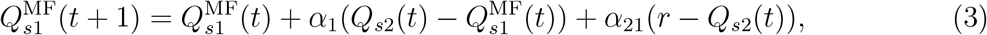

where *α*_1_ is the learning rate for the first-stage. The parameter *α*_21_ determines the extent to which the second-stage reward prediction error influences first-stage choices (which is equivalent to the temporal discounting term *α*_1_ × *λ* in the model by Daw et al. (2011)).

Model-based values were computed for each first-stage stimulus and every trial in a forward-looking manner by multiplying the state value of the best second-stage option with the state transition probabilities:

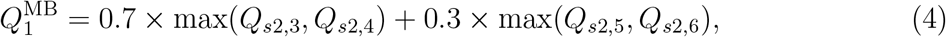

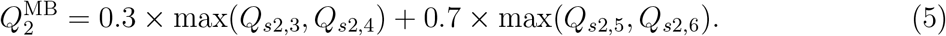

Model-based learning was simplified and the transition probabilities were not updated by explicitly modelling state prediction errors. Simulations by the authors of the original task showed that learning of state transitions quickly converge to stable values (see supplementary materials of Daw et al. (2011)).

The hybrid model computes the actual value that is used in determining first-stage choices as a weighted combination of the model-based (*Q*_MB_) and model-free (*Q*_MF_) values. The first stage Q values are computed the following way:

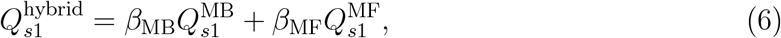

where *β*_MB_ and *β*_MF_ are the weighting parameters for model-based and model-free control, respectively (and are equivalent to *β*_1_ × *ω* and *β*_1_ × (1 − *ω*) in the model by Daw et al. (2011)). Note that in a pure model-free approach *β*_MB_ = 0, and in a pure model-based approach *β*_MF_ = 0.

Q-values for the four second-stage stimuli were updated according to the reward prediction error (Rummery and Niranjan, 1994):

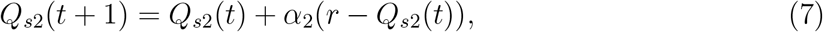

where *α*_2_ is the learning rate for the second-stage.

A first-stage choice depends on the relative difference in stimulus values between *Q*_*s*1,1_ and *Q*_*s*1,2_ and the choice *C* on the previous trial, which takes on the value 1 when the current choice equals the previous choice. The parameter *π* captures first-stage choice perseverance. Using the softmax choice function, the probability of choosing a first-stage stimulus was computed according to:

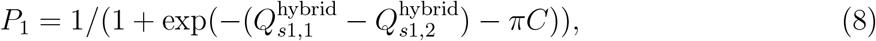

and for the second-stage:

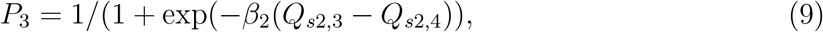

where *β*_2_ control the randomness of second-stage choices.

We used a hierarchical model-fitting strategy that takes into account the likelihood of individual participant choices given the individual participant parameters and also the likelihood of the individual participant parameters given the parameter distribution in the overall population across conditions. This two-stage hierarchical procedure is a estimation strategy of the iterative expectation-maximization algorithm (EM) (Guitart-Masip et al., 2011; MacKay, 2003). This regularises individual participants’ parameter fits, rendering them more robust toward over-fitting. To infer the maximum-a-posteriori estimate of each parameter for each participant, we set the prior distribution to the maximum-likelihood given the data of all participants and then used EM for the two conditions separately to obtain parameter estimates for each condition. The statistical significance was tested using paired t-tests with respect to the Gaussian scaled model parameters (see supplemental material for the transformation of parameters). The reported p-values were corrected for multiple comparisons using the Bonferonni method.

## Results

As expected, participants rated their subjective feelings of hunger significantly higher after 14 hours of fasting than after eating a full meal (Wilcoxon signed rank test: [*Z* = −4.84, *p <* 0.0001]), indicating that the manipulation was successful.

We first examined the influence of food deprivation on choice behaviour. During each session, a participant undertook 201 trials, of which on average 2.3 ± s.d. 3.1 trials were not completed due to failure to enter a response within the 2 seconds time limit. Food deprivation did not affect the number of missed trials [*t*_31_ = 1.19, *p* = 0.245] or median response times for first and second-stage choices (first-stage RT_hungry_ = 616 ms, RT_sated_ = 640 ms; paired t-test, [*t*_31_ = −1.29, *p* = 0.207], second-stage RT_hungry_ = 788 ms, RT_sated_ = 821 ms; paired t-test, [*t*_31_ = −1.25, *p* = 0.220]), suggesting that food deprivation did not alter overall attention in this task.

### Food deprivation only modulated model-free control

To dissociate model-based and model-free control, we measured the tendency to “stay” with the same first-stage choice as the previous trial. Model-free and model-based strategies predict distinct patterns of stay behaviour. A model-free reinforcement learning strategy predicts actions repeat when reinforced, i.e. a main effect of reward (Fig. 2A), whereas a model-based learning strategy predicts a crossover interaction between reward outcome on the second-stage and the type of transition (Fig. 2B). This arises because model-free and model-based strategies predict opposing stay probabilities on trials following a rare transition. After a rare transition, the model-free system updates the value of the first-stage *chosen* stimulus, such that reward promotes staying with the current choice. The model-based system instead updates state values, and so rewards after a rare transition will promote switching, since the *unchosen* first-stage stimulus is more likely to lead to the previously rewarded second-stage state.

**Figure 2.**
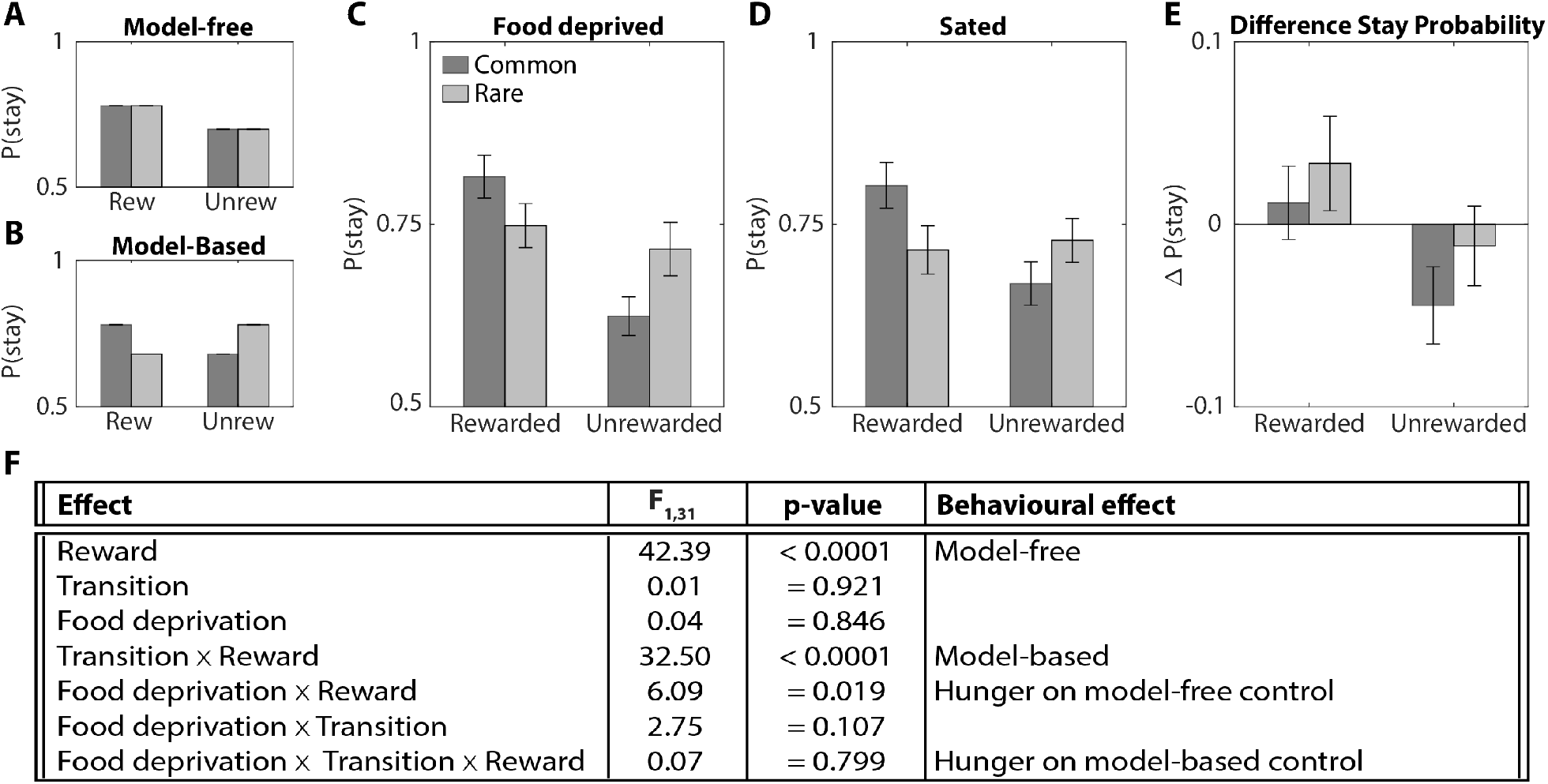
Stay-Switch behaviour at first-stage. Model-based and model-free strategies for reinforcement learning predict different patterns by which outcomes experienced after the second-stage impact first-stage choices on subsequent trials (Daw et al., 2011). **A)** Simulation of choice behaviour driven by the model-free system. Rewards increase the likelihood of choosing the same stimulus on the next trial, regardless of whether that reward occurred after a common or rare transition. **B)** Simulation of choice behaviour driven by the model-based system, which relies on an interaction between transition type and reward outcome. **C)** Experimental data of food deprived participants. **D)** Experimental data of sated participants. **E)** Change in stay probability with food deprivation. Food deprivation increased the tendency to stay after receiving a reward and increased the tendency to switch after reward omission. Error bars represent SEM. **F)** Statistics stay behaviour: three-way repeated measures ANOVA.

We examined whether the probability of staying or switching depended on the level of food deprivation (hungry or sated), the transition type (common or rare), and the reward on previous trial (reward or no reward), using a three-way repeated measures ANOVA (Fig. 2C– F). Participants used both model-free and model-based strategies to solve this task, which is consistent with previous studies (Daw et al., 2011; Deserno et al., 2015; Wunderlich et al., 2012).

The key analyses concerned whether hunger modulated these learning strategies (Fig. 2F). Food deprivation increased the tendency to repeat the same choice after receiving a reward, but not after reward omission, showing that food deprived individuals relied more on model-free strategies. Hunger did not affect overall stay behaviour or model-based strategies.

The contributions of model-free and model-based strategies to choice behaviour can be summarised with an alternative index (Eq. (1) and (2)). The current metabolic state modulated the model-free (MF) index [*t*_31_ = 2.47, *p* = 0.019], but not the model-based (MB) index ([*t*_31_ = 0.26, *p* = 0.799]; Fig. 3A). The session order and the amount of training did not alter MB or MF indices ([*p >* 0.2]; Fig. 3B). These analyses confirmed that the manipulation of metabolic state, rather than training, caused the effects observed in this study. This observation corresponds with earlier reports that extensive training did not alter the trade-off between the model-based and model-free system (Grosskurth et al., 2019).

**Figure 3.**
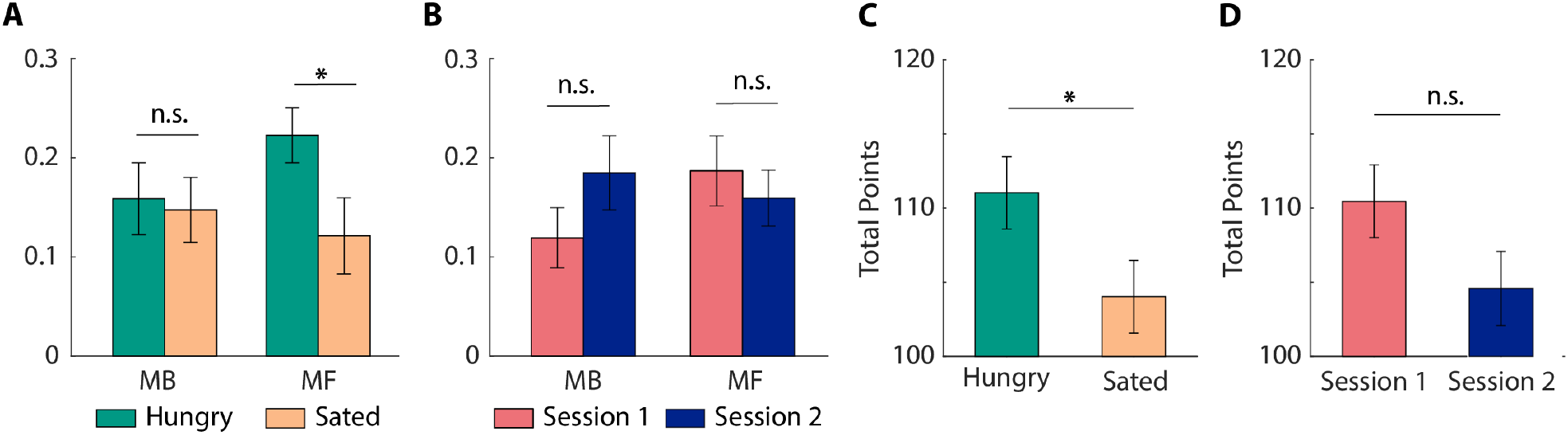
Hunger increased model-free control and performance. **A)** Food deprivation increased the relative contribution of the model-free (MF), but not model-based (MB), system to stay behaviour at stage 1. **B)** Training did not significantly alter the contribution of the MB or MF system to choice behaviour. **C)** Food deprivation increased the number of points earned. **D)** Session order had no significant impact on the number of points earned. Error bars represent SEM. \**p <* 0.05.

### Food deprivation enhanced performance

Although some studies have reported a significant correlation between overall performance and enhanced model-based control (Wunderlich et al., 2012), recent studies have shown that the number of points may not be strongly correlated to either model-based or model-free control (Akam et al., 2015; Kool et al., 2016). In this study, food deprivation enhanced performance, measured as the number of points earned in the task ([*t*_31_ = 2.25, *p* = 0.032]; Fig. 3C). The total number of points earned did not differ between the first and second testing days ([*t*_31_ = 1.84, *p* = 0.075]; Fig. 3D), suggesting that more experience with the task structure did not improve performance (Grosskurth et al., 2019).

### Computational modelling confirmed model-free effects

The behavioural pattern in Fig. 2E deviates from the pattern we expected from a pure model-free effect. Furthermore, some studies have criticised the interpretation of the stay-switch measures, in Eq. (1) and Eq. (2), as mapping onto model-free and model-based systems respectively (Grosskurth et al., 2019). Therefore, we corroborate the behavioural results with computational modelling to assess the effects of food deprivation, and attribute these effects to a specific computational process. The complexity of the task and the contribution of model-based and model-free strategies to choice behaviour were captured by a seven parameter model (Fig. 4A). These seven parameters can be grouped into four categories based on their functional roles: the model-based system, model-free system, second stage or bias. The quality of the fitting procedure was verified with a parameter recovery analysis. All parameters were well recovered (0.65 ≤ *R <* 0.95) and the model fitting procedure did not introduce spurious correlations between the other parameters (|*R*| *<* 0.4; Fig. S2).

**Figure 4.**
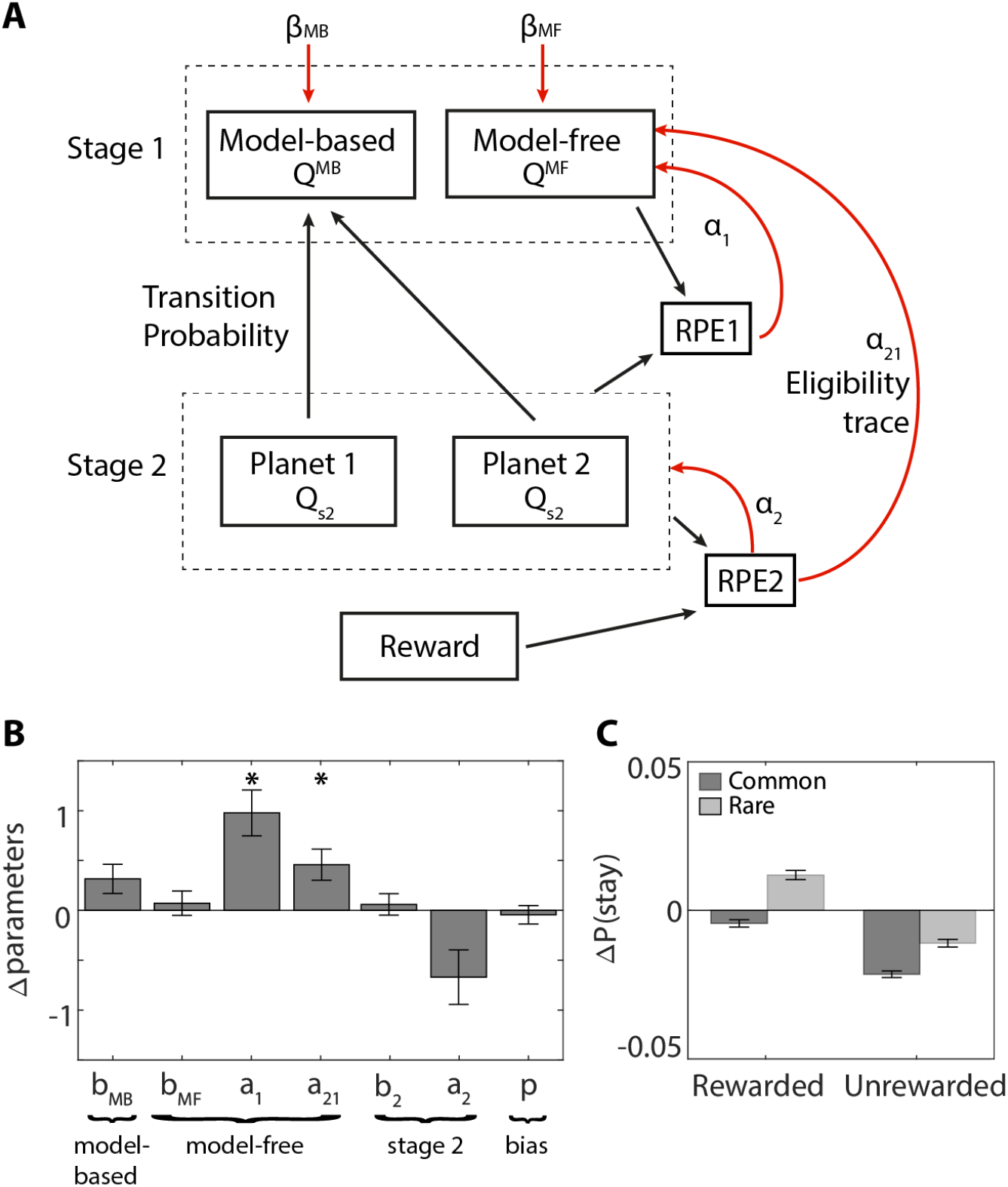
Computational modelling results. **A)** First-stage choices depend on a model-based and model-free component. Model-based values are computed by multiplying the stimulus with the highest value of both second-stage planets with their respective transition probabilities. Model-free values are updated using a first-stage reward prediction error (RPE1; scaled by *α*_1_), which captures the difference between the value of the chosen option in the first-stage and the chosen option in the second-stage (state transition), and an eligibility trace of the second-stage reward prediction error (RPE2). RPE2 captures the difference between the reward received and the value of the chosen second-stage option (scaled by *α*_2_). The model assumes that the second stage prediction error influences the first stage values with a learning rate *α*_21_, because it assumes that first stage choice leaves eligibility trace that makes *Q*^MF^ modifiable according to subsequent prediction errors. Black arrows indicate contribution to. Red arrows indicate that values are scaled by. **B)** Differences in Gaussian scaled parameters between food deprived and sated conditions. Positive values indicate that the parameter estimate was higher when food deprived compared to sated. **C)** Surrogate data simulated with the original model using best-fitting parameters for this model (Suppl. Table 1) revealed the key pattern in stay behaviour as shown by the experimental data in Fig. 2E. Error bars represent SEM. * *p <* 0.05.

We found that food deprivation significantly increased the learning rate for first-stage choices [*α*_1_, *p <* 0.001] and the contribution of second stage prediction errors [*α*_21_, *p* = 0.042]. These data suggest that hunger increased model-free learning. Hunger did not affect any of the other parameters ([*p >* 0.14]; Fig. 4B). Surrogate data generated with the best fitted parameters (Suppl. Table 1) confirmed that the key difference in stay behaviour (Fig. 2E) was captured by the estimated parameters (Fig. 4C). The modelling results confirmed that hunger increased model-free learning, without affecting model-based control, which explains the pattern observed in Fig. 2E.

## Discussion

Truly adaptive behaviour involves changing policy in response to not only outcome contingencies, but also our metabolic state. In this study, we used a sequential decision-making task to test whether hunger modulates simple reflexive decisions or prospectively planned actions. Hunger enhanced overall performance by increasing model-free control, without changing model-based control. These results indicate that hunger enhances adaptive behaviour by boosting reinforcement learning, without affecting cognitive processes that are required for future planning.

To solve the two-step task, people can either use model-free learning and simply repeat actions that yielded rewards, or they can use model-based learning to track *states* that yielded rewards, and plan actions to reach those states. These two strategies entail very different computations. These may in turn rely upon different brain systems (Lee et al., 2014) that could respond differently to resource depletion or repletion. The sequential task allowed us to track both of these modes of decision-making. In line with previous studies (Daw et al., 2011; Grosskurth et al., 2019; Wunderlich et al., 2012), our participants used a mixture of model-based and model-free strategies to solve the task.

Hunger increased the importance of model-free control, characterised by a main effect of reward and increased learning rates derived from computational modelling. When food deprived, participants won more points, suggesting that increased model-free control is beneficial to overall performance. Why might this be? To perform optimally, participants continually have to update the estimates of second-stage outcomes (aliens) according to the randomly drifting outcome probabilities. It turns out that this stochasticity imposes a low ceiling on achievable performance on this task, such that the theoretical winnings do not differ between pure model-based, model-free and random agents (Akam et al., 2015; Kool et al., 2016).

A previous study by Friedel et al. (2014) hinted towards a relationship between model-based control and metabolic need. Friedel et al. indexed the metabolic need of an individual as the change in valuation for food rewards in a separate devaluation paradigm. They observed that people who exhibited greater devaluation also showed more model-based control. In contrast, our study asks whether changes in metabolic needs have generalised effects on cognition, beyond the specific reward type that changes in value. By testing the same individuals twice on this sequential decision-making task (once in high metabolic need and once in low metabolic need), we were able to directly examine the causal effect of metabolic need on the balance between model-based and model-free control in the same individual, and correlate this with overall performance. Crucially, our observations are complementary to those of Friedel et al. (2014). Although metabolic changes did not affect model-based behaviour for *monetary* outcomes, they may still affect the value of food rewards. Indeed, the value of a particular reward increases when humans or animals are deprived of it (Aw et al., 2009; Epstein et al., 2003; Pompilio et al., 2006; Siep et al., 2009). Instead, the increased learning we observed must reflect a more general change in cognition, perhaps associated with heightened motivation.

How might such general effects arise? One mechanism may be that food deprivation acts as a mild stressor (Deroche et al., 1995). Stress itself may impair model-based control (Otto et al., 2013) and enhances reliance on model-free strategies, particularly for negative outcomes (Park et al., 2017). This may arise through the action of various metabolic hormones, as well as hypothalamic inputs to ventral prefrontal cortex. It is worth noting that the results are incompatible with hunger simply disrupting attention, which would be expected to *decrease* learning rates and performance.

Metabolic hormones mostly act on primitive, subcortical areas (Elmquist et al., 1998; Zigman et al., 2006), including the basal ganglia. These areas play an important role in model-free control (Daw et al., 2011; Deserno et al., 2015; Gläscher et al., 2010; Lee et al., 2014; Smittenaar et al., 2013), and are modulated by the current metabolic state (Abizaid et al., 2006; Aitken et al., 2016; Cone et al., 2016; Hommel et al., 2006; Papageorgiou et al., 2016). This link may support our behavioural and modelled findings that hunger increased model-free control, without affecting model-based decision-making.

Our findings may be directly relevant to populations with dysregulation of hunger. Individuals with eating disorders, but not obese individuals, exhibit increased model-free behaviour in this same task (Voon et al., 2015). Our study provides a crucial causal link, that within individual participants from an unselected population, hunger increases model-free behaviour.

To conclude, we found that increased metabolic need enhanced reflexive model-free control of behaviour, without affecting deliberate model-based control. Moreover, relying on a reflexive model-free system does not necessarily make hungry people maladaptive, when the deliberate model-based system remains unaffected.

