## Supplemental material for "Hunger improves reinforcement-driven but not planned action"

### Other reinforcement-driven behaviours were not affected

In addition to the stay-switch measure reported in Fig. 2, there are two other measures of simple reinforcement learning in this task that are independent of previous state transitions. The first measure is stay behaviour for second-stage choices, which is indicative of adaptive behaviour driven by reinforcements. The win-stay and lose-stay probabilities for second-stages were calculated for the two available stimuli together, but separately for the two planets, and divided by the number of wins or losses to give the percentage of stay behaviour for each participant. For this analysis, we assumed that the same state did not have to be visited twice in a row, because this would make the analysis dependent on the first-stage choices. Consequently, we assumed that participants were able to keep an average of the total expected value for a stimulus in their working memory. Participants were more likely to repeat their previous choice after receiving a reward (main effect of reward [$F_{1,31} = 124.50$, $p < 0.0001$]). Food deprivation did not alter overall stay behaviour (main effect of food deprivation [$F_{1,31} < 1$]) or outcome-specific stay behaviour (interaction effect of food deprivation and reward [$F_{1,31} < 1$]; Fig. S1A).

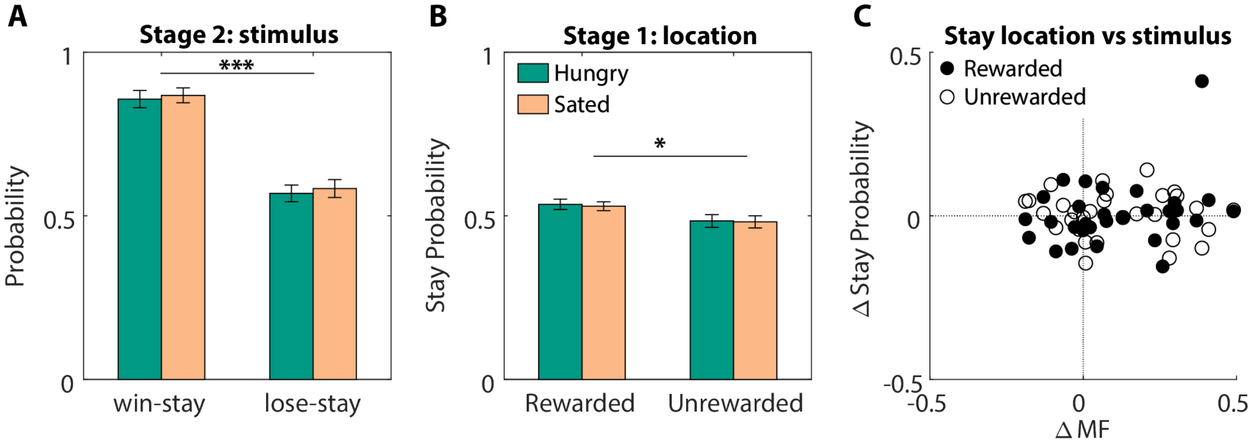


Figure S1: **Hunger did not alter other types of reinforcement learning. A)** Repetition of second-stage choices was enhanced following rewarded trials and was not altered by food deprivation. **B)** Repetition of first-stage actions was enhanced following rewarded trials, regardless of stimulus identity or the level of food deprivation. **C)** There was no correlation between first-stage action repetition and the model-free (MF) values of first-stage choices. Action repetition did not contribute to model-free control of stage 1 choices. Error bars represent SEM. * $p < 0.05$, *** $p < 0.001$.

The second measure is stay-switch behaviour for outcome-irrelevant information (Shahar et al., 2019). Participants may associate actions (i.e. choosing left or right), rather than stimulus identity, with reward outcomes and repeat an action after receiving a reward on the previous trial for that action. Given that only the stimulus identity, not the stimulus location, has predictive effects, this type of behaviour could be seen as a measure of aberrant incidental learning, in which a value is assigned to an action that was not inherently rewarding. Participants repeated the same action more often after receiving a reward (main effect of reward [$F_{1,31} = 6.34$, $p = 0.017$]). Food deprivation did not alter overall stay behaviour (main effect of food deprivation [$F_{1,31} = 0.19$, $p = 0.664$]) or outcome-specific stay behaviour (interaction effect of food deprivation and reward [$F_{1,31} < 1$]; Fig. S1B). The significant effect of reward on action repetition did not affect the contribution of the model-free system to first-stage choices (Fig. S1C). Together, these analyses show that food deprivation selectively affected model-free control of first-stage choices, but not all types of reinforcement-driven learning.

### Computational modelling

#### Parameter transformations

Before fitting the parameters, we applied logistic/exponential transformations to transform bounded parameters into normal distributed parameter values $x_i \sim \mathcal{N}(\mu_x, \sigma_x)$, with a population mean of $\mu_x$ and a standard deviation of $\sigma_x$. We transformed [0,1]- bounded $\alpha$ into a Gaussian scale using the logistic function:

$$\alpha = 1/(1 + \exp(-a)), \tag{1}$$

and the logarithmically scaled $\beta$ and $\pi$ using the exponential function:

$$\beta = \exp(b) \tag{2}$$

We denote the model parameters by Greek letters and the Gaussian transformation by their respective latin letters. Normally distributed parameters allow for the use of parametric tests to identify differences between conditions.

#### Parameter recovery

To validate the parameter estimates generated by the fitting procedure, we conducted a parameter recovery analysis. For each parameter, we generated samples from the marginalised posterior distribution of the fit to get realistic parameters that could describe choice behaviour. The generated parameters were uncorrelated ($|R| < 0.3$), allowing for testing whether the fitting procedure introduced any confounding factors. Using these parameters, we simulated choice behaviour for 32 virtual subjects following the task design described in the method section. We then used a hierarchical model fitting procedure to estimate parameters for the simulated data ("Recovered parameters"). We assessed the quality of the parameter recovery by comparing the true parameters used to simulate data with the recovered parameters. We calculated the Pearson correlation between all pairs of recovered parameters to test whether the fitting procedure introduced spurious correlations. A strong correlation between the true and recovered parameters indicates a good recovery of the parameters and reliable model fitting results. All parameters were well recovered ($0.65 < R < 0.95$) and the model fitting procedure did not introduce spurious correlations between the other parameters ($|R| < 0.4$; Fig. S2).

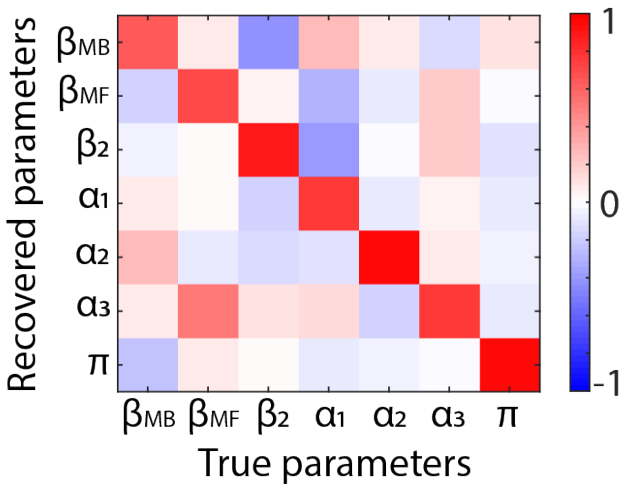


Figure S2: **Parameter recovery.** Correlation matrix of the free parameters used to generate the simulated data ('True parameters') and the obtained parameters by applying the parameter estimation procedure on the simulated data ('Recovered parameters'). Bright red values indicate a strong correlation between the true and recovered parameter value and therefore a good parameter recovery.

#### Surrogate data

To confirm that our model can capture our key behavioural findings, we generated data for 32 virtual subjects on this task using the individual best-fitting parameters (Suppl. Table 1) and the same trajectories of reward probabilities as experienced by the participant. Each simulation was repeated 30 times to obtain an average. These data were then subjected to a stay-switch analysis. We found an identical pattern of effects in these generated data as observed empirically in our participants (Fig. 4C).

| | $\beta_{\mathbf{MB}}$ | $\beta_{\mathbf{MF}}$ | $\beta_2$ | $\alpha_1$ | $\alpha_2$ | $\alpha_{21}$ | $\pi$ |
|---|---|---|---|---|---|---|---|
| **Sated** | | | | | | | |
| 25th percentile | 1.43 | 1.40 | 2.13 | 0.15 | 0.30 | 0.08 | 0.30 |
| Median | 2.25 | 2.10 | 3.64 | 0.23 | 0.49 | 0.14 | 0.67 |
| 75th percentile | 4.22 | 2.96 | 5.15 | 0.30 | 0.64 | 0.23 | 1.38 |
| **Food deprived** | | | | | | | |
| 25th percentile | 1.65 | 1.86 | 2.37 | 0.34 | 0.11 | 0.17 | 0.27 |
| Median | 3.09 | 2.35 | 4.23 | 0.45 | 0.40 | 0.22 | 0.72 |
| 75th percentile | 7.03 | 3.01 | 4.93 | 0.54 | 0.58 | 0.26 | 1.13 |

Table 1: **Best-fitting model parameters estimates.** Separately shown for the sated and food deprived condition as median and quartiles across participants.
